# Beta-Hydroxybutyrate Inhibits Bronchial Smooth Muscle Contraction

**DOI:** 10.1101/2025.02.24.639075

**Authors:** V. Amanda Fastiggi, Madeleine M. Mank, Matthew A. Caporizzo, Matthew E. Poynter

## Abstract

Asthma is a chronic respiratory condition characterized by airway inflammation, remodeling, and hyperresponsiveness to triggers causing airway constriction. Bronchial smooth muscle plays a critical role by narrowing airways, leading to obstruction and breathing difficulties, often exacerbated by mast cell infiltration and histamine release. Whereas current treatments, including bronchodilators, corticosteroids, and biologics provide effective management for most patients, alternative therapies are needed for difficult-to-treat asthma. Recent research highlights the potential of therapeutic ketosis, achieved through dietary interventions or supplementation with exogenous ketones, to reduce airway hyperresponsiveness and inflammation. Ketone bodies, known for providing energy during carbohydrate scarcity, also influence asthma by activating cell-surface receptors and transporters. *In vivo*, interventions like weight loss and caloric restriction increase ketone body levels, correlating with improved asthma symptoms, reduced oxidative stress, and inflammation. These effects suggest ketone bodies, particularly β-hydroxybutyrate, may play a therapeutic role in mitigating bronchoconstriction and smooth muscle contraction in asthma. We utilize human bronchial smooth muscle cells (*in vitro*) and mouse precision-cut lung slices (PCLS) (*ex vivo*) to assess the effects of BHB on histamine-induced bronchoconstriction. Brightfield microscopy showed that BHB reduces contraction in human bronchial smooth muscle cells, an effect involving free fatty acid receptor 3 (FFAR3) activation. Light microscopy of PCLS revealed that BHB inhibits airway narrowing and cellular extrusion, demonstrating its ability to mitigate bronchoconstriction by suppressing smooth muscle contraction. These results implicate bronchial smooth muscle as a cellular target of therapeutic ketosis, an important contributor to the beneficial effects of BHB in preclinical models of asthma.

**Graphical Abstract:** In accordance with journal requirements, a graphical abstract will be submitted if we are invited to revise this manuscript.

## Introduction

Asthma is a chronic, heterogenous, and widespread respiratory syndrome affecting patients’ breathing that is characterized by hyperresponsiveness to broncho-constrictive triggers, inflammation, and remodeling of the airway, and with which more than a quarter of a billion people are diagnosed worldwide^1^. Airway hyperresponsiveness can be assessed through the methacholine challenge test, which serves as a diagnostic tool to evaluate airway smooth muscle function and aid in the clinical diagnosis of asthma^2,3^. Bronchial smooth muscle plays a key role in the pathophysiology of asthma by causing airway narrowing through cellular contraction, leading to obstruction and difficulty breathing^4^. Stimulation of bronchial smooth muscle can also contribute to lung inflammation through the production of pro-inflammatory cytokines^5^.

Bronchial smooth muscle can be both affected by the inflammatory environment as well as contribute to it. Mast cells can infiltrate into the smooth muscle tissue in allergic asthma and thereby induce or augment airway hyperresponsiveness^6^. This infiltration can cause bronchial smooth muscle cells to release pro-inflammatory chemotactic cytokines (chemokines)^7,8^, and the mast cells release mediators such as histamine^8,9^ which contributes to airway obstruction by causing smooth muscle contraction, increasing bronchial secretions, and provoking mucosal edema^9^. Bronchoconstriction was amongst the first biological effects described for histamine^10^ and can cause bronchial smooth muscle contraction to the same extent as M1 muscarinic receptor agonists (*e.g.*, methacholine) and has been suggested to generate more contraction in peripheral tissue^11^.

Current treatments for asthma include broncho-relaxing β-agonists, anti-inflammatory corticosteroids, and biological immunotherapies targeting the innate and adaptive immune response^12^. While most patients achieve effective disease control, some with ‘difficult-to-treat’ asthma require alternative or additional therapies. Our lab^13,14^ and others^15^ have reported the efficacy of augmenting circulating ketone body concentrations, known as “therapeutic ketosis”, to mitigate the pathophysiological manifestations in mouse models of obese and allergic asthma. Ketone bodies, β-hydroxybutyrate (BHB) and acetoacetate (AcAc), are endogenously produced in the liver from fatty acids^16,17^, either through diet^17,18^ or adipose tissue mobilization during energy demand^17^, and are then circulated to cells throughout the body.

Ketone bodies have been implicated in modulating key pathological processes in asthma. Originally described as an energetic substrate for the production of ATP in the Krebs Cycle during time of carbohydrate scarcity^19^, ketone bodies have also been reported to exert their effects through the stimulation of cell-surface receptors, such as G-protein coupled receptors HCAR2 (GPR109a) and FFAR3 (GPR41) ^15,17,20–23^, or by uptake through transporters such MCT1^22,24^. Additionally, ketone bodies act as antioxidants^25,26^ and have anti-inflammatory properties, including the suppression of nuclear factor-κB (NF-κB) activation^25,25^ and inhibition of the NLRP3 inflammasome, which reduces IL-1β production^27–29^. *In vivo*, dietary interventions such as weight loss and alternate-day caloric restriction raise BHB levels, which correlate with reduced asthmatic symptoms, including lower oxidative stress and inflammation in obese asthmatics^30–32^.

Notably, ketone augmentation or therapeutic ketosis is well-tolerated in human subjects^36,37^. Early ketone body elevations are associated with decreased asthma symptoms in obese asthmatic subjects following bariatric surgery^33,34^, undergoing alternate-day caloric restriction^30^, and during treatment with GLP-1R agonists^35^. Therapeutic ketosis can also be achieved through providing exogenous ketones or ketogenic precursors (*e.g.*, ketone esters). Therapeutic ketosis is achieved through feeding a high fat/ low carbohydrate diet or by supplementing the normal diet with ketones esters, significantly and substantially decreases asthma associated methacholine responsiveness^13,14^ and also decreases methacholine responsiveness in non-asthmatic mice^13^. As methacholine functions through the activation of bronchial smooth muscle^38,39–41^, we sought to study these cells more directly to explore the impact of ketone bodies on their activity. We developed a reliable *in vitro* model using human bronchial smooth muscle cells (HBSMC), enabling direct observation of BHB’s effects on morphological change in this cell type^14^. Herein, we used HBSMC as an *in vitro* model of bronchial smooth muscle along with precision cut lung slices (PCLS) as an *ex vivo* model to more thoroughly assess the effects of ketone bodies on these cells and the mechanisms whereby they function.

PCLS are a valuable *ex vivo* model for studying airway reactivity and contraction, closely mimicking lung architecture and cellular diversity^42–45^. They bridge the gap between *in vitro* and *in vivo* studies and are particularly useful in asthma research, demonstrating altered responses like hyperresponsiveness and bronchoconstriction when exposed to various agonists^42,45–47^. Building on the utility of PCLS in asthma research, this study explores the potential of ketone bodies, particularly BHB, to directly influence bronchial smooth muscle contraction, addressing gaps in our understanding of their therapeutic potential.

Although connections between the underlying mechanisms of asthma and the benefits of ketone body augmentation are increasing, further evaluations are still needed, specifically studies on the capacity of ketone bodies to affect bronchial smooth muscle directly^13,14^. We hypothesized that BHB can mitigate bronchoconstriction by inhibiting contraction of bronchial smooth muscle. Our objectives were to assess the effectiveness of BHB in reducing histamine-induced bronchial smooth muscle contraction *in vitro* and *ex vivo* to identify the mechanisms by which BHB may influence these effects. Further understanding of the efficacy and mechanisms of BHB attenuation of contraction in bronchial smooth muscle could provide insight into novel targets for the treatment of asthma and asthmatic symptoms.

## Materials and Methods

### Study Approval

The animal experiments were reviewed and approved by the University of Vermont’s Institutional Animal Care and Use Committee (PROTO202000195), in accordance with the recommendations in the *Guide for the Care and Use of Laboratory Animals*, prepared by the Institute of Laboratory Animal Resources, National Research Council, and published by the National Academy Press (revised 2011). Studies involving potentially hazardous materials were reviewed and approved by the University of Vermont’s Institutional Biosafety Committee (REG201900052).

### Human Bronchial Smooth Muscle Cell Culture

Primary human bronchial smooth muscle cells (HBSMC) isolated from a 45-yr-old female patient with asthma (Lonza, Morristown, NJ, Lot No. 00194850, Batch No. 0000195154) were cultured in smooth muscle cell growth medium-02 BulletKit (Lonza, CC-3182) according to the manufacturer’s instructions at 37°C in 95% humidified air containing 5% CO_2_. The cells were used within the first seven passages to ensure proper smooth muscle phenotype. Cell authentication was performed by Lonza (negative Factor VIII-related antigen, positive α-Actin expression) and cells tested negative for mycoplasma (MycoDect Mycoplasma Detection Kit (Alstem, Richmond, CA)) before being utilized for experiments.

### Microscopy-Based Contraction Assay

For simultaneous exposure and stimulation experiments, human bronchial smooth muscle cells (HBSMC) were plated at 5×10^4^ cells/cm^2^ in 1mL of media in a 12-well plate and allowed to grow for 24 hours at 37°C and 5% CO_2_ to ensure sub-confluency for better visualization. HBSMC were exposed for 5 minutes with vehicle or 1.25-10 mM beta-hydroxybutyric acid (BHBA) (Sigma-Aldrich, St. Louis, MO, Cat No.166898), sodium beta-hydroxybutyrate (NaBHB) (Sigma Aldrich, Cat No.54965), (R)-beta-hydroxybutyrate ((R)-BHBA) (Sigma Aldrich, Cat No.54920), (S)-beta-hydroxybutyrate ((S)-BHBA) (Sigma Aldrich, Cat No.54925)), FFAR3 agonist AR420626 (AR) (Caymen, Ann Arbor, MI, Cat No. 17531), or a combination of MCT1 inhibitor AZD3965 (AZD) (MedChemExpress, Monmouth Junction, NJ, Cat No. HY12750) and BHBA, before being stimulated with 0.1–10 mM histamine (Sigma Aldrich, Cat No.H7125).

In pre-exposure experiments with subsequent stimulation, HBSMC were plated at 5×10^4^ cells/cm^2^ in 1mL of media in a 12-well plate and allowed to grow for 24 hours at 37°C and 5% CO_2_. After the initial 24 hours, cells were exposed to vehicle or 1.25-10 mM BHBA, NaBHB, (R)-BHBA, or (S)-BHBA for another 24 hours before being washed and stimulated with 2.5 mM histamine.

Cells were imaged with an EVOSxl system (Thermo Fisher, Waltham, MA) to acquire brightfield microscopy images at 20X magnification captured at 10 second intervals for 5 minutes. For experiments, through time-lapse brightfield microscopy, the contraction of individual cells was identified by visual cellular shape change and the reduction of intercellular space. Cellular area changes were quantified using the polygon tool in ImageJ (FIJI) to outline the perimeter of individual cells. The enclosed area was measured with ImageJ software both before and after agonist stimulation. The pre-stimulation value was normalized to 100%, and percent change was calculated. Percent contraction is presented as the reciprocal of the percent change, in which a negative percent change represents a decrease in cellular area, indicating contraction. Each individual cell analyzed represents n=1.

### Precision Cut Lung Slices (PCLS)

6–12-week-old female and male C57BL/6J mice were euthanized, and their whole lungs were inflated with 40°C 1.5% low-melting point agarose in phosphate buffered saline (PBS) through the cannulated trachea and cooled in ice-cold PBS. The right murine lobe was uniformly cut into 150μM thick slices using a 7000smz-2 Vibratome (Campden Instruments, Lafayette, IN) maintained at 4°C, at a speed of 0.7mm/s. The freshly cut PCLS were placed in sterile, room temperature 1X PBS for imaging or into submerged tissue culture plates for culture with DMEM-F12 medium supplemented with 1X penicillin/streptomycin, 1μg/mL insulin, and 1X Primocin for continued incubation of the PCLS in 5% CO_2_ at 37°C. Cultured PCLS were imaged within 4 days.

### Ex Vivo Live Tissue Imaging

For simultaneous exposure and stimulation experiments, PCLS were exposed for 10 minutes with vehicle or 10mM BHBA or NaBHB while images were acquired and were subsequently stimulated with 10mM histamine. In pre-exposure with subsequent stimulation experiments, PCLS were exposed to 10mM BHBA or NaBHB for 24 hours at 37°C and 5% CO_2_, and then placed in PBS to be imaged and stimulated with 10mM histamine. Live tissue contraction of the PCLS was imaged using the Zeiss Airyscan 2 confocal microscope imaging system in conjunction with the ImageJ Micro-Manager 2.0.0 Multi-Dimensional Acquisition (FIJI). Images were acquired at a frame rate of 1 frame/second and were analyzed using ImageJ (FIJI) software measuring relative diameter change to calculate percent airway narrowing. A 3D-printed PLA wafer was used during imaging to weigh down the PCLS to increase efficiency and reproducibility of imaging. Each individual airway represents n=1.

### Statistical Analyses

All experiments included multiple biological replicates for each condition. Outliers were identified and removed using the ROUT method (Q=1%) and the cleaned data were analyzed using unpaired one-way ANOVA with post-hoc multiple comparisons: Tukey’s test (for comparing all means) or Dunnett’s test (for comparing each mean to a control), were performed using GraphPad Prism 10.2.3 (GraphPad Software, Inc., La Jolla, CA). Data are presented as means ± SEM from representative experiments. P values below 0.05 are considered statistically significant, and significance levels are indicated in the figure legends.

## Results

### Histamine induces human bronchial smooth muscle cell contraction

Histamine is well-documented as a potent inducer of airway smooth muscle contraction that causes bronchoconstriction^10^. Using human bronchial smooth muscle cells (HBSMC) as an *in vitro* model, we confirmed that histamine provokes a dose-dependent contraction of these cells (**Figure 1A**). The contraction induced by 2.5 mM histamine was visualized using light microscopy, with comparative images captured before stimulation and 5 minutes after stimulation (**Figure 1B**). Post-stimulation images clearly show contracted cells, highlighted by red arrows, emphasizing the effect of histamine exposure. Quantitative analysis revealed that histamine elicited significant contraction compared to the vehicle control. Additionally, dose-dependent increases in contraction were statistically significant across all comparisons, except for the transition from 1 mM to 2.5 mM histamine. These results demonstrate the robust and measurable effect of histamine on bronchial smooth muscle contraction, enabling the examination of ketone bodies on this asthma-related phenotype.

**Figure 1.**
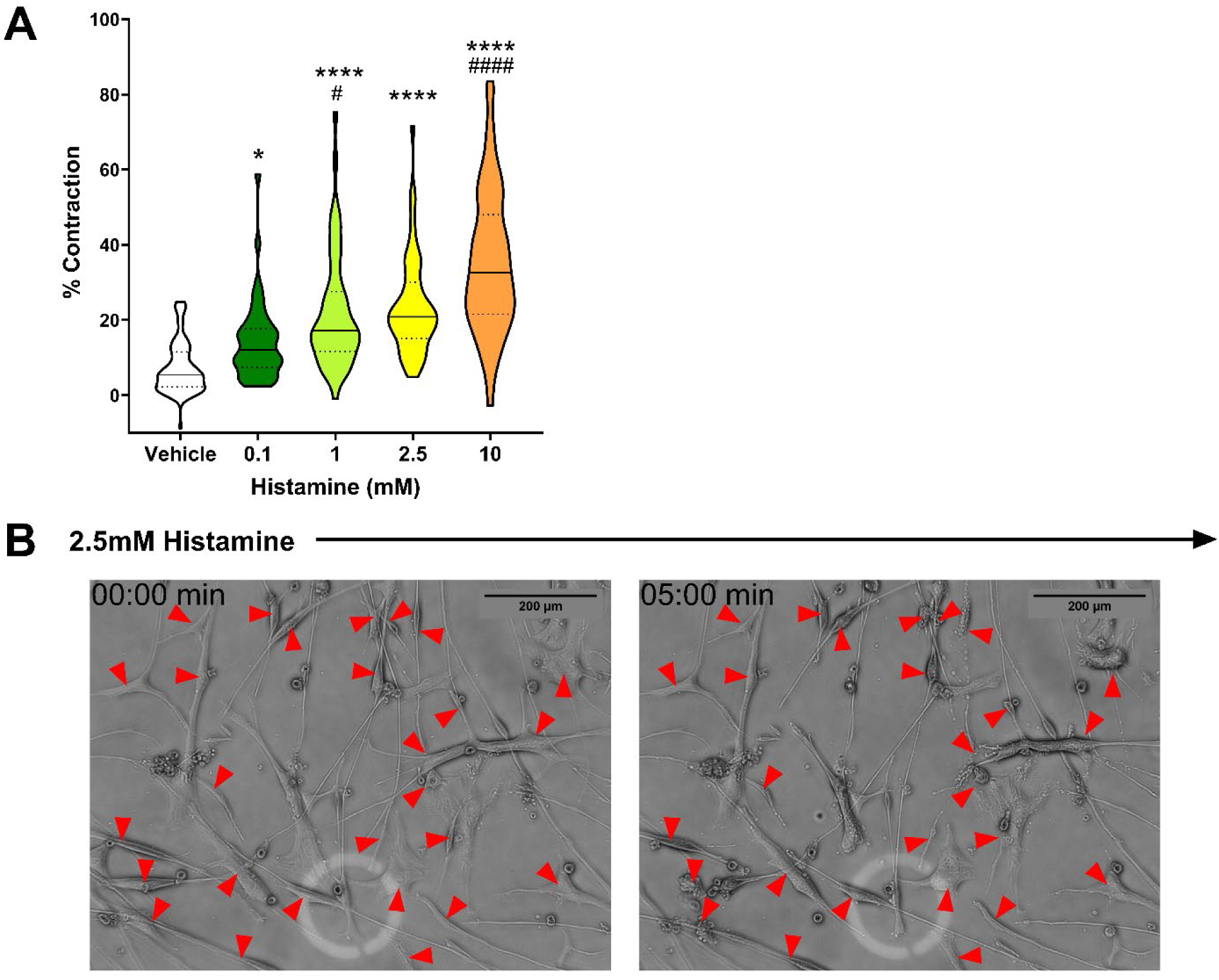
Histamine dose-dependently provokes HBSMC contraction. HBSMCs were sequentially stimulated *in vitro* with increasing concentrations of histamine for 5 minutes each and percent contraction of individual cells was quantitated. n=44-89 per group; the values presented represent a normalized dataset created by combining results from three individual studies. ∗*P* ≤ 0.05, ∗∗*P* ≤ 0.01, ∗∗∗*P* ≤ 0.001, ∗∗∗∗*P* ≤ 0.0001 compared to the vehicle, #*P* ≤ 0.05 ####*P* ≤ 0.0001 compared to the previous concentration of histamine **(A)**. Visualization of contracting cells through sample movie **(Supplemental Video 1)** stills of 2.5mM histamine response of HBSMC (minutes: seconds), with red arrows pointing to individual contracting cells (scale bars, 200μm) **(B)**.

### Beta-hydroxybutyrate (BHB) attenuates histamine induced human bronchial smooth muscle cell contraction

In our recent studies, we demonstrated the therapeutic benefits of elevating circulating ketone bodies—a state known as therapeutic ketosis—in mouse models of both obese^13^ and allergic asthma^14^. This intervention effectively mitigated methacholine-induced airway hyperresponsiveness and significantly improved lung function *in vivo*^13,14^. Building on these findings, we investigated the effects of ketone bodies on histamine-induced contraction in human bronchial smooth muscle cells (HBSMC) using our validated *in vitro* system. Whereas exposure to 2.5 mM histamine elicited pronounced HBSMC contraction, cells that were co-treated with 2.5 mM beta-hydroxybutyric acid (BHBA), (R)-beta-hydroxybutyric acid ((R)-BHBA), (S)-beta-hydroxybutyric acid ((S)-BHBA), or sodium beta-hydroxybutyrate (NaBHB) exhibited significantly attenuated histamine-induced contraction (**Figure 2**). Notably, none of the BHB forms tested induced contraction in the absence of histamine, indicating their specific role in modulating histamine-evoked responses.

**Figure 2.**
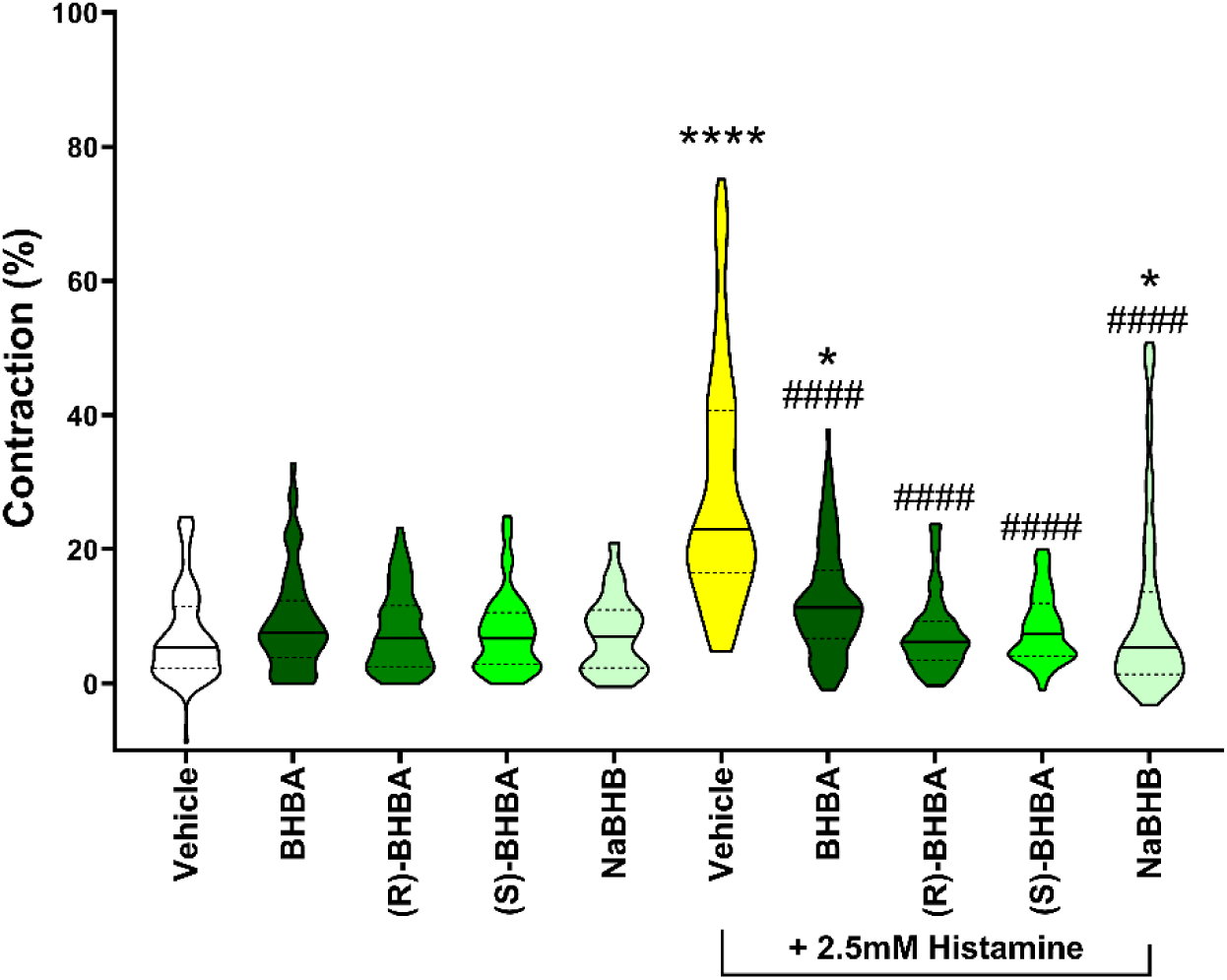
BHB decreases histamine-induced HBSMC contraction. HBSMCs were simultaneously stimulated *in vitro* for 5 minutes with 2.5mM histamine in the presence of 2.5mM BHBA, (R)-BHBA, (S)-BHBA, or NaBHB and percent contraction of individual cells was quantitated. n=24-100 per group; the values presented represent a normalized dataset created by combining results from three individual studies. Supplemental material of representative movies for each condition can be provided upon request. ∗∗∗∗*P* ≤ 0.0001 compared to the vehicle, ####*P* ≤ 0.0001 compared to histamine.

Interestingly, the attenuation of contraction was more pronounced with the racemic BHBA and the individual enantiomers (R)-BHBA and (S)-BHBA compared to sodium beta-hydroxybutyrate (NaBHB). These findings suggest that the pH of the beta-hydroxybutyrate compound contributes to its efficacy, highlighting the potential for targeted therapeutic strategies leveraging specific BHB formulations.

### Beta-hydroxybutyric acid dose-dependently inhibits histamine-induced HBSMC contraction

Building on the observed inhibitory effects of various forms of BHB on histamine-induced contraction, we next investigated whether these effects were dose-dependent. Human bronchial smooth muscle cells (HBSMC) were simultaneously stimulated with 2.5 mM histamine and exposed to increasing, biologically relevant concentrations of beta-hydroxybutyric acid (BHBA) for 5 minutes *in vitro*. By quantifying the percentage of cell contraction, we found that BHBA effectively attenuated histamine-induced contraction in a dose-dependent manner (**Figure 3**). Notably, all tested concentrations of BHBA significantly reduced contraction compared to the histamine control, further supporting its potential to modulate airway smooth muscle reactivity.

**Figure 3.**
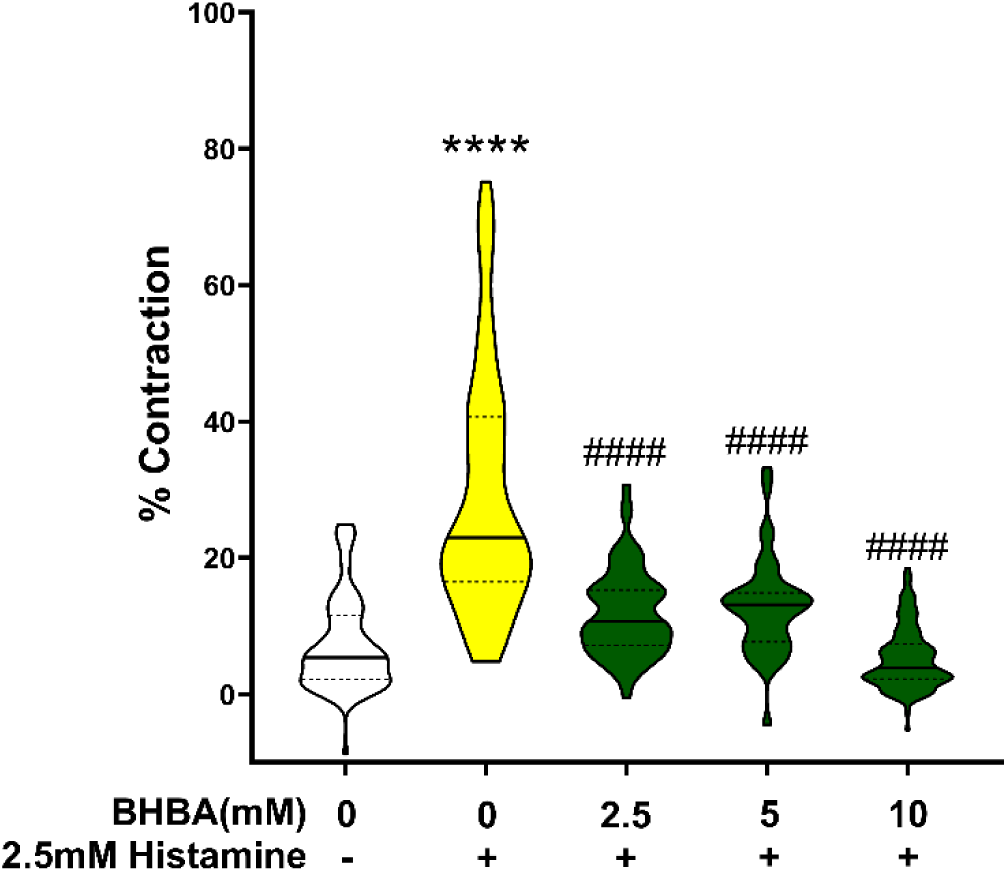
BHBA dose-dependently inhibits histamine-induced HBSMC contraction. HBSMCs were simultaneously stimulated *in vitro* for 5 minutes with 2.5mM histamine in the presence of increasing concentrations of beta-hydroxybutyric acid (BHBA) and percent contraction of individual cells was quantitated. n=40-114 per group; the values presented represent a normalized dataset created by combining results from three individual studies. ∗∗∗∗*P* ≤ 0.0001 compared to the vehicle, ####*P* ≤ 0.0001 compared to histamine.

### Pre-exposure of HBSMC to BHBA attenuates histamine-induced contraction

As our previously reported *in vivo* studies have modeled endogenous ketone augmentation through dietary interventions that provide elevated systemic concentrations of these molecules over a protracted period, we sought to replicate this exposure in an *in vitro* system. To mimic endogenous ketone elevation, human bronchial smooth muscle cells (HBSMC) were pre-exposed to biologically relevant concentrations of beta-hydroxybutyric acid (BHBA) for 24 hours, followed by washing and subsequent stimulation with histamine. Pre-exposure to BHBA significantly attenuated histamine-induced contraction in a dose-dependent manner (**Figure 4**), consistent with the inhibitory effects observed during simultaneous exposure of BHBA and histamine. Notably, all pre-exposure concentrations demonstrated significant reductions in contraction compared to the histamine control, which received an appropriate vehicle pre-exposure. Interestingly, when comparing the two exposure methods, the simultaneous exposure of BHBA with histamine proved more effective at reducing contraction than the pre-exposure approach.

**Figure 4.**
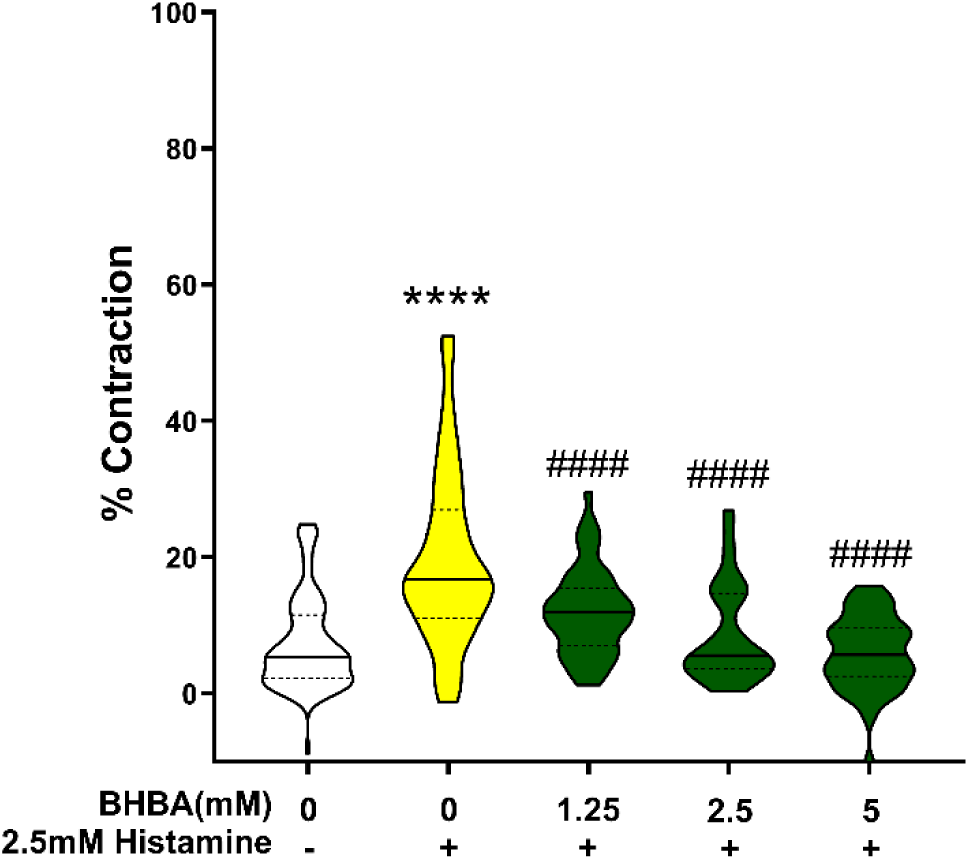
BHBA pretreatment attenuates histamine-induced HBSMC contraction. HBSMCs were untreated (Vehicle) or exposed *in vitro* to increasing concentrations of beta-hydroxybutyric acid (BHBA) for 24 hours, washed, and then stimulated with 2.5mM histamine for 5 minutes and percent contraction of individual cells was quantitated. n=40-89 per group; the values presented represent a normalized dataset created by combining results from three individual studies. ∗∗∗∗*P* ≤ 0.0001 compared to the vehicle, ####*P* ≤ 0.0001 compared to histamine.

### Pre-exposure to BHB or a Free Fatty Acid Receptor (FFAR)3 agonist inhibits histamine-induced HBSMC contraction

Given that beta-hydroxybutyrate (BHB) has been suggested to act as a ligand for FFAR3 and exert its beneficial effects through an FFAR3-dependent pathway^21–23,48^, we investigated whether activating FFAR3 could replicate the inhibitory effects of BHB on histamine-induced contraction in HBSMC. To test this possibility, cells were pre-exposed to biologically relevant concentrations of BHB to model endogenous ketone augmentation, as well as to the FFAR3 agonist AR420626 (AR). Both pre-exposure to BHB and AR significantly attenuated histamine-induced HBSMC contraction (**Figure 5**), suggesting that the observed inhibitory effects of BHB may indeed be mediated, at least in part, through the activation of FFAR3.

**Figure 5.**
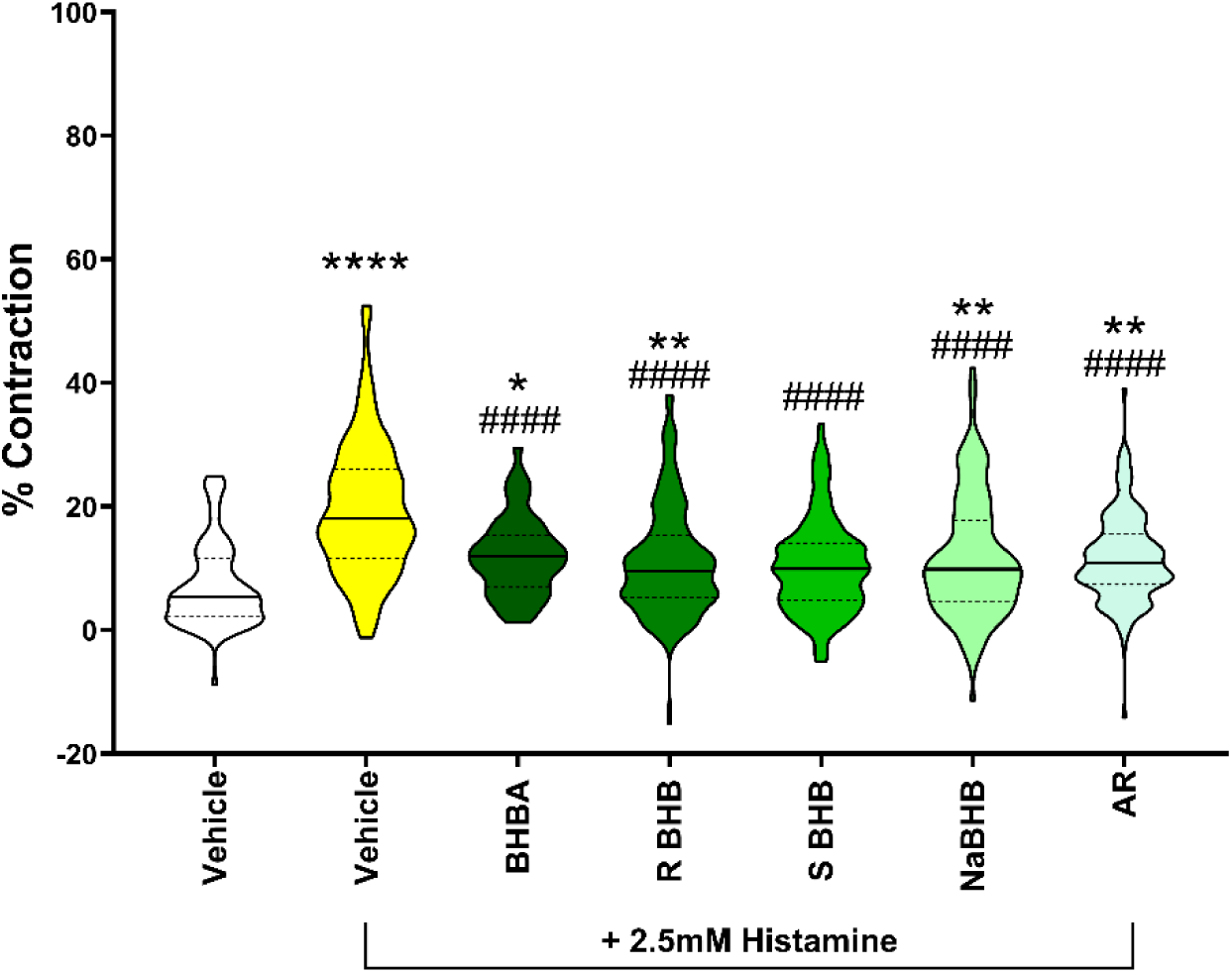
Pretreatment with BHB compounds or FFAR3 agonist attenuates histamine-induced HBSMC contraction. HBSMCs were untreated (Vehicle) or exposed *in vitro* with beta-hydroxybutyric acid (BHBA), (R)-BHBA, (S)-BHBA, NaBHB, or FFAR3 agonist (AR) for 24 hours, washed, and then stimulated with 2.5mM histamine for 5 minutes and percent contraction of individual cells was quantitated. n=51-216 per group; the values presented represent a normalized dataset created by combining results from three individual studies. Supplemental material of representative movies for each condition can be provided upon request. ∗*P* ≤ 0.05, ∗∗*P* ≤ 0.01, ∗∗∗*P* ≤ 0.001, ∗∗∗∗*P* ≤ 0.0001 compared to the vehicle, ####*P* ≤ 0.0001 compared to histamine.

### FFAR3 activation is sufficient to attenuate histamine-induced HBSMC contraction

The ability of the FFAR3 agonist AR420626 (AR) to attenuate histamine-induced contraction in a simultaneous stimulation mirrors the effects observed with pre-exposure to BHBA, (R)-BHBA, (S)-BHBA, or NaBHB. This similarity suggests that these compounds may exert their inhibitory effects through a shared mechanism or pathway. In our model of exogenous ketone augmentation, simultaneous stimulation with histamine in the absence or presence of BHB compounds or AR once again attenuated histamine-induced HBSMC contraction (**Figure 6**).

**Figure 6.**
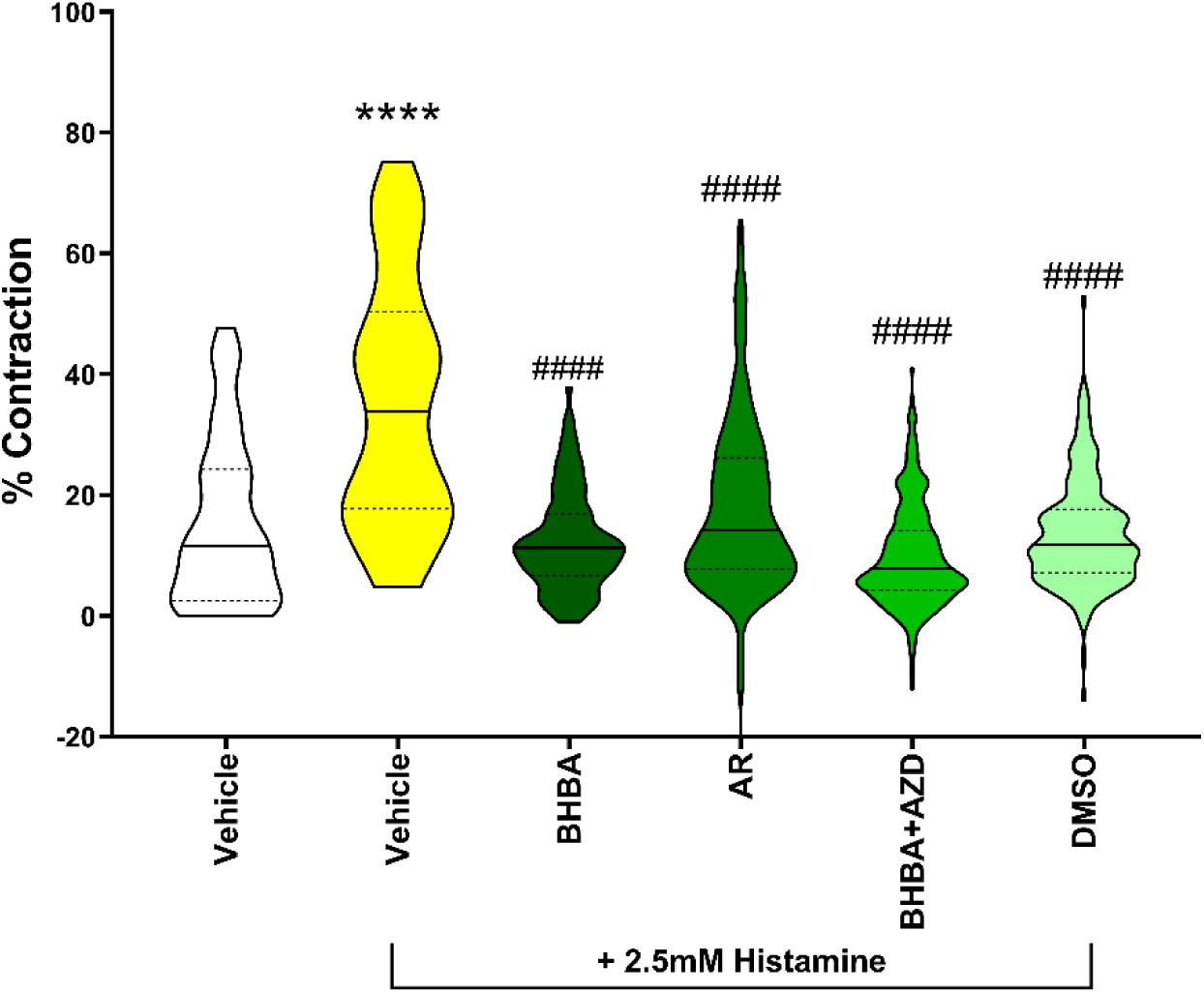
FFAR3 activation is sufficient to inhibit histamine-induced HBSMC contraction. HBSMCs were untreated (Vehicle) or exposed *in vitro* with 2.5mM histamine in the presence of 2.5mM beta-hydroxybutyric acid (BHBA), 50µM FFAR3 agonist (AR), or a combination of 40nM MCT1 inhibitor (AZD) and 2.5mM BHBA for 5 minutes and percent contraction of individual cells was quantitated. DMSO was used as a vehicle control for the AR and AZD groups. n= 42-291 per group; values are inclusive of studies performed three times. Supplemental material of representative movies for each condition can be provided upon request.∗∗∗∗*P* ≤ 0.0001 compared to the vehicle, ####*P* ≤ 0.0001 compared to histamine.

Ketone bodies, particularly BHB, have been proposed to mediate their beneficial effects via uptake through Monocarboxylic Transporter 1 (MCT1) (21, 37). To evaluate the involvement of this pathway, we tested the MCT1 inhibitor AZD3965 (AZD) in the presence of the BHB compounds. Notably, the presence of AZD did not block the inhibitory effects of BHB on histamine-induced contraction, indicating that these effects are independent of MCT1-mediated uptake. Interestingly, the use of DMSO as a vehicle for both AR and AZD also resulted in unexpected inhibition of histamine-induced contraction. This observation underscores the need for further investigation into the potential off-target effects of DMSO in this context.

### Histamine induces airway narrowing in murine PCLS

It has been reported that in murine models of allergic asthma, methacholine (McH) exposure causes pronounced bronchoconstriction of mouse lungs that were immune-primed with either ovalbumin (OVA) or house dust mite (HDM)^14,49^ and that treatment of human and guinea pig PCLS with histamine causes a decrease in airway lumen area^50^. It was also recently reported that agonist-induced bronchoconstriction in the airways can cause pathological airway epithelial crowding, triggering cellular extrusion into the airway lumen^49^. Using live tissue imaging of murine PCLS, we observed that histamine causes dose-dependent narrowing of the airway by causing cellular extrusion during bronchoconstriction (**Figure 7A**). The cellular extrusion induced by 10 mM histamine was visualized using light microscopy, with comparative images captured before stimulation and 10 minutes after stimulation (**Figure 7B**). Post-stimulation images clearly show extruded cells, highlighted by red arrows, emphasizing the effect of histamine exposure. Quantitative analysis revealed that histamine elicited significant airway extrusion compared to the vehicle control (**Figure 7C**). Interestingly, we did not observe the same robust airway narrowing in our sequential dose-response studies in which the 10mM histamine condition only induced a mean value of about 5% airway narrowing, whereas in the studies in which a single stimulation of 10mM histamine was used, a mean value of about 55% airway narrowing or closure was observed.

**Figure 7.**
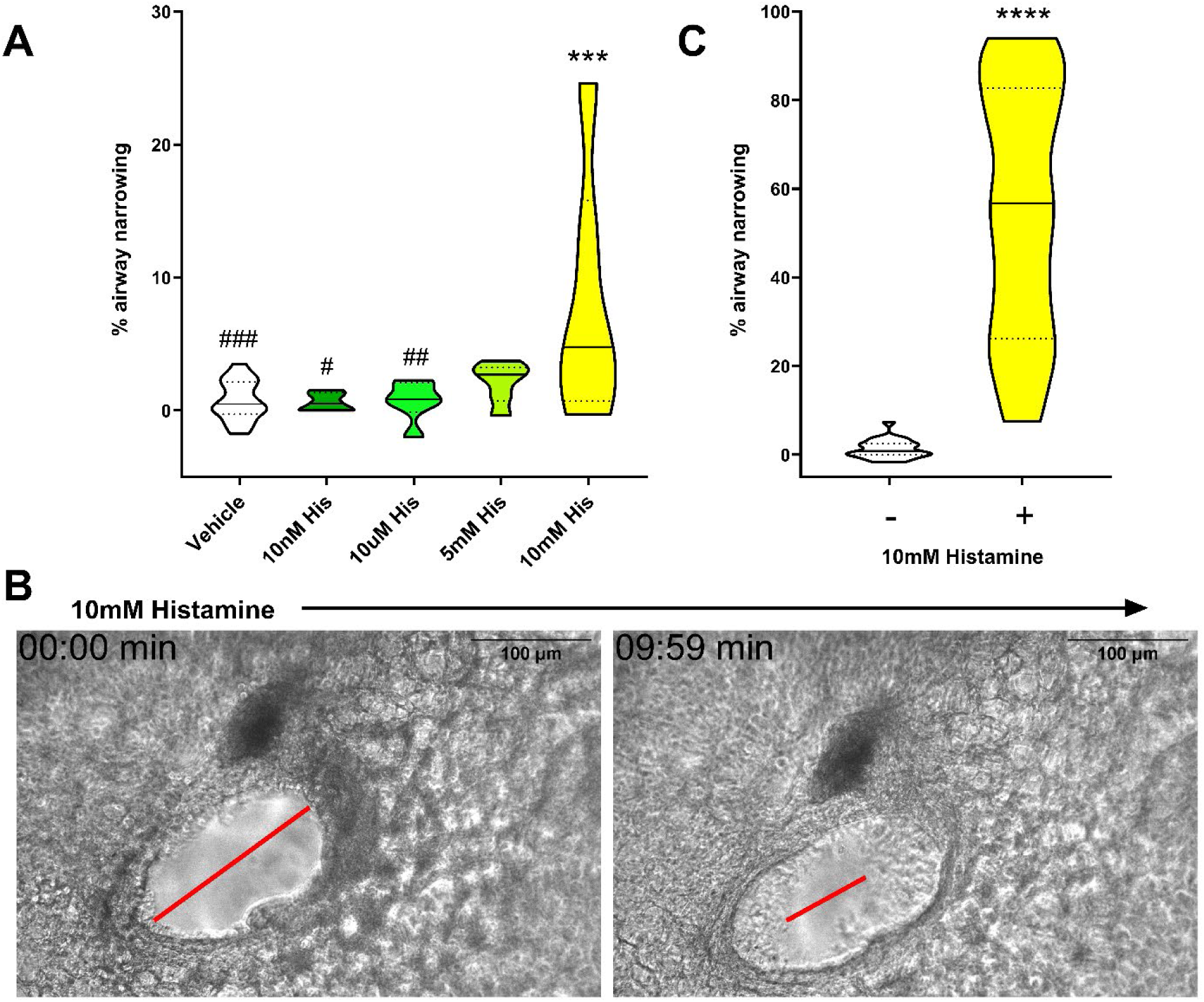
Histamine induces cellular extrusion in murine PCLS. Murine PCLS were sequentially stimulated *ex vivo* with increasing concentrations of histamine for 10 minutes each and percent airway narrowing of individual airways was quantitated. n=6 per group; the values presented represent a normalized dataset created by combining results from three individual studies. ∗∗∗*P* ≤ 0.001 compared to the vehicle, #*P* ≤ 0.05, ##*P* ≤ 0.01, ###*P* ≤ 0.001 compared to 10mM histamine **(A)**. Visualization of cellular extrusion through sample movie **(Supplemental Video 2)** stills of 2.5mM histamine response of PCLS (minutes: seconds), with red arrows pointing to cellular extrusions (scale bars, 100µm) **(B)**. Quantification of percent airway narrowing PCLS stimulated with 10mM histamine. n=27-30 per group; the values presented represent a normalized dataset created by combining results from three individual studies. ∗∗∗\**P* ≤ 0.0001compared to the vehicle **(C)**.

### Exogenous BHB stimulation is sufficient to inhibit airway cellular extrusion in PCLS

We have previously reported that BHB has inhibitory effects on allergen-induced pro-inflammatory cytokine secretion^14^ in human bronchial epithelial cells, providing some evidence that BHB already has effects directly on this relevant cell type. Using *in vitro* model systems, histamine can disrupt barrier integrity through adverse effects on tight junction integrity^51–53^ and fluid hypersecretion^52,54^. Building on our findings *in vitro* of the inhibitory effects of BHB on histamine-induced HBSMC contraction, we investigated the effects of ketone bodies on histamine-induced airway narrowing *ex vivo*.

Exposure to 10 mM histamine elicited a pronounced phenotype of cellular extrusion and a significant increase in the percentage of airway narrowing occurring, as expected (**Figure 8A**). However, in PCLS exposed to 10 mM beta-hydroxybutyric acid (BHBA), or sodium beta-hydroxybutyrate (NaBHB), histamine-induced airway narrowing, and cellular extrusion visualized using light microscopy, was significantly attenuated, with comparative images captured before stimulation and 10 minutes after stimulation (**Figure 8B**) in the presence of BHBA or NaBHB. Post-stimulation images clearly show extruded cells, highlighted by red arrows, emphasizing the effect of BHB exposure compared to the histamine positive control.

**Figure 8.**
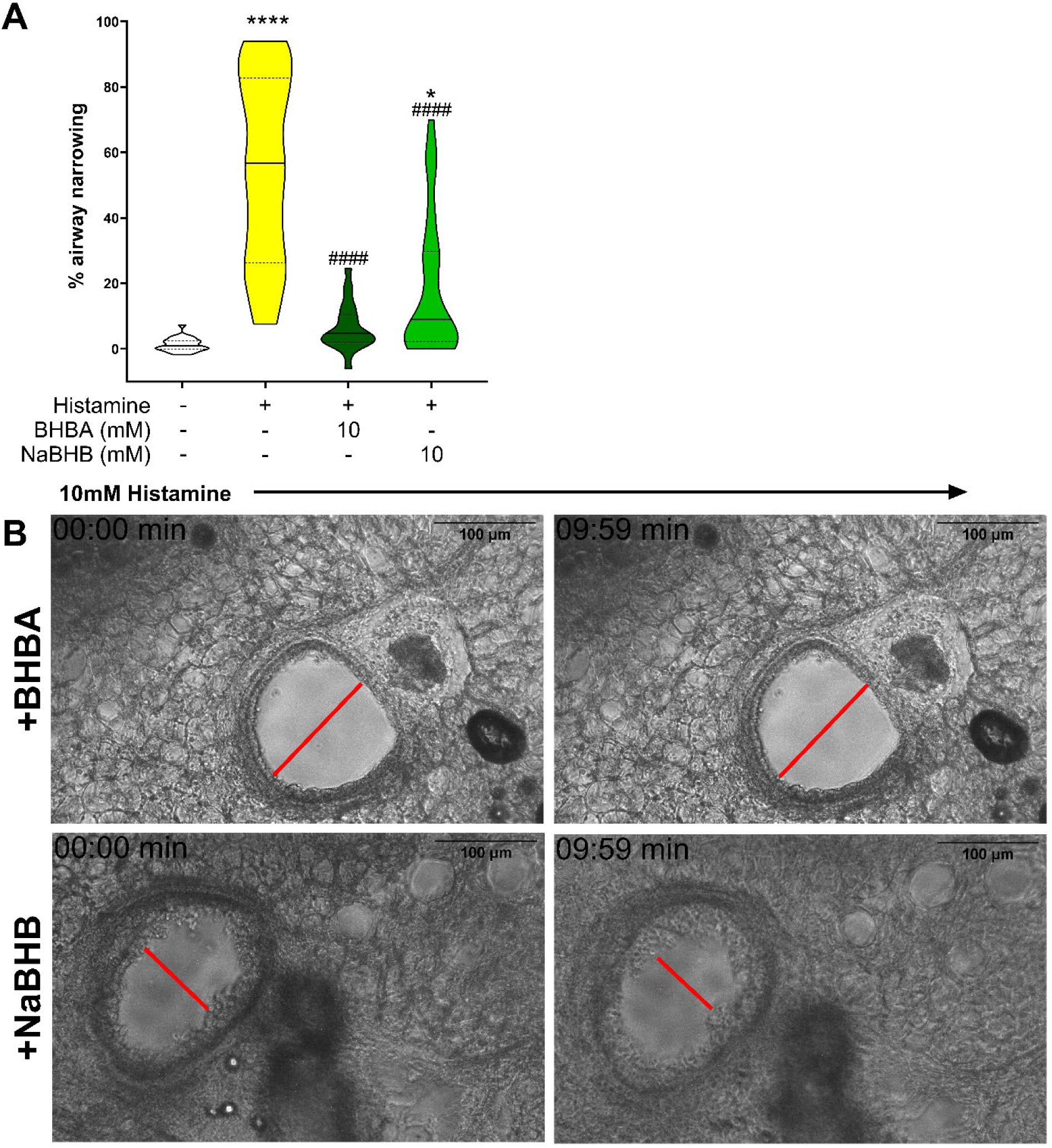
BHB attenuates histamine-induced cellular extrusion in PCLS. Murine PCLS were simultaneously stimulated *ex vivo* with 10mM histamine for 10 minutes each in the presence of 10mM BHBA or NaBHB and percent airway narrowing of individual airways was quantitated. and percent airway narrowing of individual airways was quantitated. n=22-32 per group; the values presented represent a normalized dataset created by combining results from three individual studies. ∗*P* ≤ 0.05, ∗∗∗∗*P* ≤ 0.0001 compared to the vehicle, ####*P* ≤ 0.0001 compared to 10mM histamine **(A)**. Visualization of cellular extrusion through sample movie stills of the 10mM histamine response of PCLS in the presence of BHBA **(Supplemental Video 3)** and NaBHB **(Supplemental Video 4)** (minutes: seconds), with red arrows pointing to cellular extrusions (scale bars, 100µM) **(B)**.

### Pre-exposure of PCLS with BHB attenuates histamine-induced cellular extrusion

As our previously reported *in vivo* studies have modeled endogenous ketone augmentation through dietary interventions, we sought to replicate this exposure in an *ex vivo* system. To model endogenous ketone elevation over a protracted period, PCLS were pre-exposed to biologically relevant concentrations of 10mM beta-hydroxybutyric acid (BHBA) or sodium beta-hydroxybutyrate (NaBHB) for 24 hours, followed by subsequent stimulation with histamine in the imaging medium containing no BHB. Pre-exposure to BHBA significantly attenuated histamine-induced cellular extrusion, consistent with the inhibitory effects observed during simultaneous exposure of BHBA and histamine. Notably, all pre-exposure conditions demonstrated reductions in cellular extrusion, although the effect of NaBHB only trended towards inhibition and was not statistically significant compared to the histamine control that received an appropriate vehicle pre-exposure (**Figure 9A**). Interestingly, when comparing the two methods of exposure, the simultaneous exposure of BHBA with histamine proved more effective at reducing cellular extrusion than the pre-exposure. Post-stimulation images clearly show extruded cells, highlighted by red arrows, emphasizing the effect of BHB exposure compared to the histamine positive control (**Figure 9B**).

**Figure 9.**
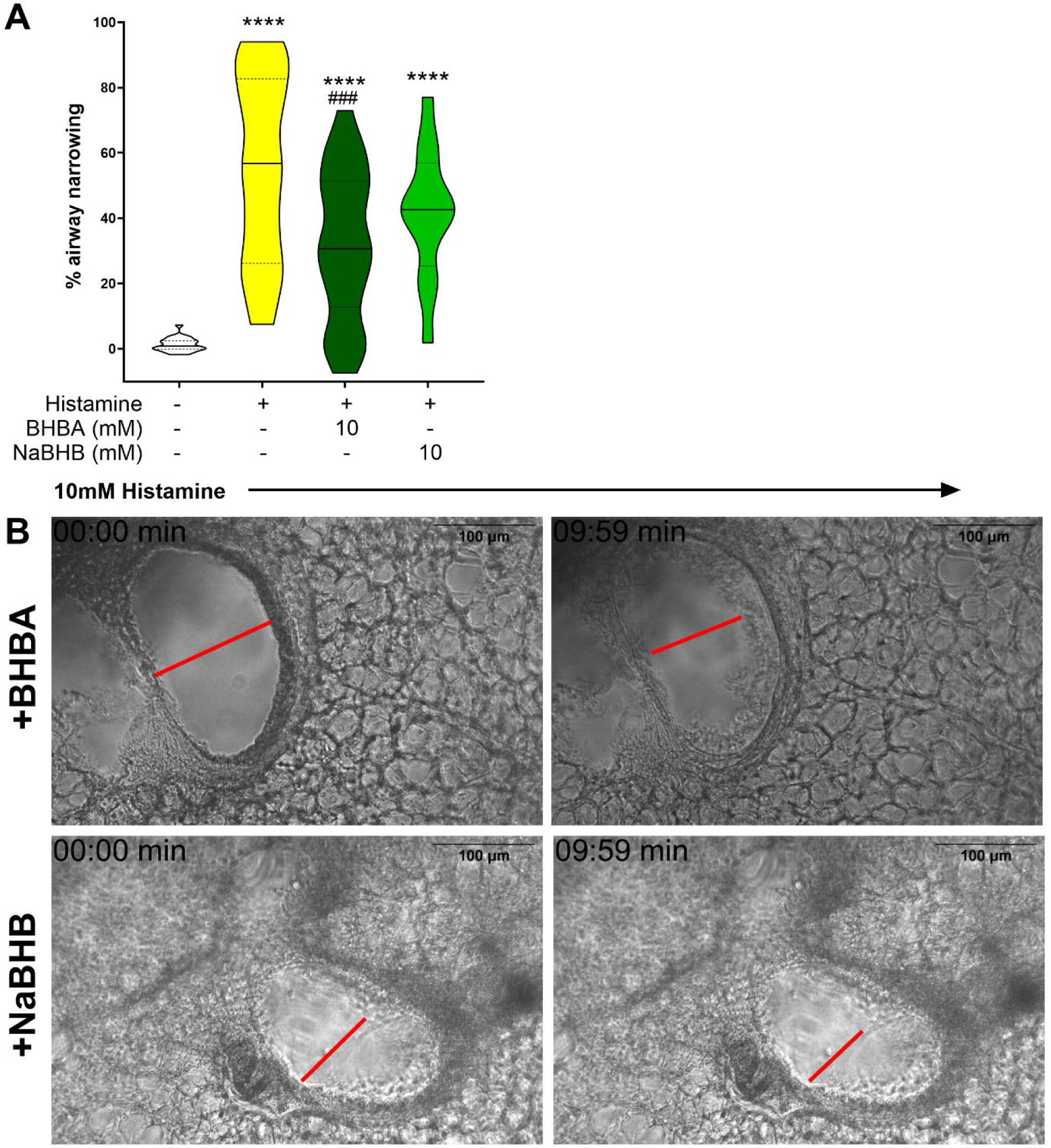
Pre-exposure of PCLS to BHB attenuates histamine-induced cellular extrusion. Murine PCLS were untreated (Vehicle) or exposed *ex vivo* with beta-hydroxybutyric acid (BHBA), or NaBHB for 24 hours, washed, and then stimulated with 10mM histamine for 10 minutes each for 10 minutes in the presence of 10mM BHBA or NaBHB and percent airway narrowing of individual airways was quantitated. n=12-23 per group; the values presented represent a normalized dataset created by combining results from three individual studies. ∗*P* ≤ 0.05, ∗∗∗∗*P* ≤ 0.0001 compared to the vehicle, ####*P* ≤ 0.0001 compared to 10mM histamine Visualization of cellular extrusion through sample movie stills of 10mM histamine response of PCLS when pre-exposed to BHBA **(Supplemental Video 5)** and NaBHB **(Supplemental Video 6)** (minutes: seconds), with red arrows pointing to cellular extrusions (scale bars, 100µM) **(B)**.

## Discussion

The increasingly prevalent asthma epidemic requires novel alternative or additional therapies to treat severe or ‘difficult-to-treat’ endotypes and improve quality of life for those with this chronic and heterogeneous syndrome. Therapeutic ketosis is one such approach that has been gaining attention as a potential therapy due to its many beneficial effects in a myriad of pathological conditions and diseases^55–59^. Pertaining to asthma, therapeutic ketosis achieved through dietary interventions—such as a ketogenic diet, ketogenic precursor supplementation, or ketone ester intake—reduces methacholine hyperresponsiveness, a key pathophysiological feature of preclinical obese associated and allergic-associated asthma models^13,14^ and inhibits pathological activities of several associated cells ^13–15^. In models of asthma, distal airway and peripheral lung dysfunction occur^60^, and our previous work has shown that augmented BHB concentrations provide beneficial effects on both^13,14^. Ketone body augmentation decreased markers of methacholine hyperresponsiveness, airway resistance, tissue damping, and tissue elastance, physiological variables which are markedly increased by heterogeneous ventilation to the distal airways commonly due to variations in airway narrowing^60,61^, and sensitive to contraction of the peripheral airways. Therefore, we speculated that the mechanism through which ketones attenuate airway hyperresponsiveness may directly involve the bronchial smooth muscle. As reported herein, our studies demonstrate that BHB inhibits bronchial smooth muscle contraction.

The mechanisms by which BHB can elicit these effects remain uncertain but may include activation of various cell surface receptors such as free fatty acid receptor 3 (FFAR3)^17,22,23^, or through modulation of intracellular signaling regulating cellular contraction and extrusion. Given that BHB has been proposed to demonstrate efficacy across a wide range of applications and activation of many G-protein coupled receptors (GCPRs)^15,16,20–23^, perhaps including the induction of tachyphylaxis, it likely operates through a fundamental mechanism that is universally relevant to these processes such as ion exchange^22,29,62^ through calcium signaling^22,23^ or adjusting membrane potential^62^.

Our *in vitro* studies using human bronchial smooth muscle cells (HBSMCs) demonstrate that histamine-induced contraction is attenuated by BHB, including its racemic mixture and the individual enantiomers (R)-BHB and (S)-BHB. Since only (R)-BHB can be efficiently metabolized by β-hydroxybutyrate dehydrogenase to form acetyl-CoA that is subsequently converted into ATP and used as an energy substrate ^63^, we confirmed that the attenuation of contraction occurs independently of BHB’s metabolic role. Additionally, the MCT1 inhibitor AZD3965 failed to block BHB’s effects, indicating that MCT1-mediated uptake is not involved.

We further explored the influence of pH by comparing sodium BHB (NaBHB, pH 9.73) and beta-hydroxybutyric acid (BHBA, pH 2.09). When diluted to biologically relevant concentrations in buffered media, both BHBA and NaBHB attenuated histamine-induced contraction *in vitro*, although BHBA was more effective suggesting that while BHB itself is sufficient, acidity enhances its effect. This mechanism may also explain differences in histamine-induced cellular extrusion *ex vivo*. Additionally, the FFAR3 agonist AR420626 produced similar inhibitory effects, implying that FFAR3 activation may mediate the inhibitory effects of BHB. Confirmation would require an FFAR3 antagonist, although a compound with this activity is not yet commercially available. Nevertheless, these findings provide the rationale for future studies, such as examining the effects of BHB in FFAR3 knockdown cells and FFAR3 knockout mice.

The simultaneous exposure to BHBA with histamine was more effective at reducing histamine-induced HBSMC contraction and PCLS cell extrusion than the BHBA pre-exposure approach. However, it is likely that the application of ‘therapeutic ketosis’ in a clinical setting would provide consistently elevated BHB concentrations that could be modeled *in vitro* and *ex vivo* using the pre-exposure approach without removing the BHB before assessment of histamine-induced contraction and cell extrusion.

Cellular extrusion occurs when the epithelial cell lining the airway becomes too crowded and start to extrude into the airway lumen^49,65^. In previous work in this field, it was determined that pathological crowding due to bronchoconstriction causes epithelial cells to extrude into the airway lumen resulting in inflammation and mucus secretion in models of both human and mouse PCLS^49^. Unlike our mouse PCLS model, the previously mentioned studies used an HDM-primed allergic asthma model in which bronchoconstriction was induced with methacholine. In contrast, our model did not involve an *in vivo* allergic asthma setup but instead used histamine as an immune-relevant agonist to induce bronchoconstriction *ex vivo*. Notably, we found that histamine elicited the same cellular extrusion effects as those observed in the HDM-primed mouse PCLS stimulated with methacholine. Additional future studies include augmenting ketone body concentrations *in vivo* in preclinical asthma models and determining whether the attenuation of histamine-induced effects on PCLS are retained *ex vivo*.

There are several limitations to our findings. In our previous *in vivo* murine studies, methacholine was used as an agonist to assess the asthma-associated lung function phenotype of methacholine hyperresponsiveness, whereas in our *in vitro* and *ex vivo* studies, we utilized histamine as an airway smooth muscle contraction-inducing agonist. Histamine contributes to asthma by triggering inflammation and airway constriction^6,8,9^. During an asthma attack, histamine is released from mast cells as part of an IgE-mediated immune response to allergens^64^ or via their IgE-independent stimulation by irritants^9^, making it a more relevant agonist for asthma studies. In contrast, methacholine is a synthetic compound modeling endogenous acetylcholine that is used in clinical diagnostics to induce airway constriction without triggering the inflammatory response characteristic of asthma^2^. Thus, histamine effectively models relevant mechanisms of asthma exacerbations and is appropriate for the studies reported herein.

There are several limitations to our findings. Namely, we conducted *in vitro* studies using primary human bronchial smooth muscle cells and *ex vivo* studies with mouse precision cut lung slices (PCLS) instead of human subjects. While informative, human cell studies fail to capture the prolonged and complex nature of human asthma. They offer a reductionist approach by focusing on a single cell type and a specific agonist. In our previous *in vivo* murine studies, methacholine was used as an agonist to assess the asthma-associated lung function phenotype of methacholine hyperresponsiveness, whereas in our *in vitro* and *ex vivo* studies, we utilized histamine as an airway smooth muscle contraction-inducing agonist. Histamine contributes to asthma by triggering inflammation and airway constriction^6,8,9^. During an asthma attack, histamine is released from mast cells as part of an IgE-mediated immune response to allergens^64^ or via their IgE-independent stimulation by irritants^9^, making it a more relevant agonist for asthma studies. In contrast, methacholine is a synthetic compound modeling endogenous acetylcholine that is used in clinical diagnostics to induce airway constriction without triggering the inflammatory response characteristic of asthma^2^. Thus, histamine effectively models relevant mechanisms of asthma exacerbations and is appropriate for the studies reported herein.

Therapeutic ketosis is being applied to respiratory diseases in the clinical setting, including asthma^66,67^ and Cystic fibrosis^68^. Whereas the asthma trials are providing medium-chain triglyceride supplementation as a substrate for ketone body formation *in vivo*, the Cystic fibrosis trial is providing the ketone ester precursor that more rapidly and efficiently boosts circulating ketone body concentrations and is the same compound we have employed in mouse models of obese asthma and allergic asthma^13,14^. The inclusion of therapeutic ketosis in clinical trials reflects growing scientific interest and recognition of its potential. Providing ketone ester supplementation in asthmatic subjects merits future study. Positive outcomes from these studies would provide robust support for incorporating therapeutic ketosis into treatment protocols, encouraging further exploration of its use as a complementary or alternative asthma therapy. This ongoing research underscores the significance of studying ketone-based interventions, emphasizing the need for continued investigation into their mechanisms and therapeutic potential.

## Abbreviations

AcAc: acetoacetate
ANOVA: analysis of variance
BHB: beta-hydroxybutyrate
BHBA: beta-hydroxybutyric acid
FFAR3: free fatty acid receptor 3
HBSMC: human bronchial smooth muscle cells
HCAR2: hydroxycarboxylic acid receptor 2
NaBHB: sodium beta-hydroxybutyrate
PBS: phosphate-buffered saline
R-BHB: (R)-beta-hydroxybutyrate
S-BHB: (S)-beta-hydroxybutyrate
SEM: standard error of the mean

